# Beneficial substrate partitioning boosts non-aqueous catalysis in de novo enzyme-alginate beads

**DOI:** 10.1101/2021.04.12.439416

**Authors:** Richard Stenner, H. Adrian Bunzel, Adrian J. Mulholland, J. L. Ross Anderson

**Affiliations:** School of Biochemistry, University of Bristol, University Walk, BS8 1TD Bristol, UK; Bristol Centre for Functional Nanomaterials, HH Wills Physics Laboratory, University of Bristol, BS8 1TL Bristol, UK; Centre for Computational Chemistry, School of Chemistry, University of Bristol, BS8 1TS Bristol, UK

## Abstract

Synthetic reactions often require solvents incompatible with biocatalysts. Here, we encapsulate a *de novo* heme-containing enzyme, C45, in calcium-alginate hydrogel beads to facilitate heterogeneous biocatalysis in neat organic solvents. Post-encapsulation, C45 retains activity even when the beads are suspended in organic solvents. In particular, the carbene transferase activity of C45 is enhanced when reactions are performed in aprotic, non-polar solvents such as hexane and toluene. Activity-solvent dependencies reveal that this activity boost is likely due to beneficial partitioning of the substrate into the beads from the organic phase. Furthermore, encapsulation facilitates enzyme recovery and recycling after the reaction. Such encapsulation opens up novel opportunities for biocatalysis in organic solvent systems, combining desired solvent properties of organic chemistry with enzymatic selectivity and proficiency.

---

Catalysis is key to the economical and sustainable synthesis of industrial chemicals.^[1]^ Growing concerns over feedstock security, rising energy costs, and sustainability have amplified demands for greener catalysts. To satisfy these demands, biocatalysis has emerged as an attractive, green alternative to synthetic processes driven by traditional, and often toxic, small molecule catalysts. Furthermore, biocatalysts often offer significantly enhanced activities and stereoselectivities over abiological syntheses. Consequently, the biocatalytic toolbox is undergoing constant expansion through natural enzyme discovery,^[2]^ creation of *de novo* biocatalysts,^[3]^ and directed evolution of promiscuous activities in natural and *de novo* enzymes.^[4]^ Such bioengineering endeavors have afforded new biocatalysts with tailored selectivities, capable of accepting non-natural substrates, and even catalyzing completely abiological transformations.^[4]^

Although tailor-made enzymes can potentially catalyze countless organic transformations, they are limited to operating under predominantly aqueous conditions to maintain their folded state; this precludes the targeting of organic transformations where substrates have poor aqueous solubilities.^[5]^ Except for some enzymes (e.g. lipases and selected extremophilic enzymes) that retain activity in high concentrations of organic solvents,^[6]^ most biocatalysts are either unstable or insoluble in non-aqueous solvents, resulting in diminished or undetectable activity.^[7]^ However, from a biotechnological perspective, there are clear advantages for non-aqueous enzymology including: increased solubility of organic compounds; suppression of unwanted side reactions; ease of product extraction; and enzyme recoverability. To achieve this, it has been established that protein immobilization can shield enzymes from solvent-induced denaturation by providing a protective aqueous layer.^[8]^ In particular, encapsulation in calcium-alginate gels provides a simple method to facilitate such organic enzymology.^[9]^ Alginate is a polysaccharide in the cell walls of brown algae that instantaneously forms gelatinous water-insoluble beads upon dropwise addition to aqueous CaCl2 solutions. This approach has proven advantageous in the past, giving rise to potent biotechnological methodologies facilitating enzyme recycling and the assembly of flow reactors.^[9f–j]^ However, enzyme encapsulation often decreases reaction yields when compared to free enzyme-catalyzed transformations.^[9d, 9e]^ The origins of that activity loss are not well understood, limiting biocatalytic applications.

Here, we examine the interplay of alginate encapsulation and activity of a catalytically promiscuous *de novo* enzyme in organic solvents. We recently created C45, a *de novo* thermostable α-helical heme-containing enzyme. C45 is a proficient peroxidase^[10]^ that can also act as a highly selective carbene transferase, catalyzing cyclopropanation, N-H insertion, carbonyl olefination, and homologous ring expansion reactions.^[11]^ Here, we show that immobilizing C45 in calcium-alginate beads not only protects the *de novo* enzyme from neat organic solvents, but also boosts the reaction yields for carbene insertion reactions; this is achieved through beneficial partitioning of the substrates between beads and solvent (Fig. 1). This highlights a general and somewhat overlooked approach to perform proficient, selective, and abiotic enzymatic catalysis in non-aqueous solvent systems.

**Figure 1:**
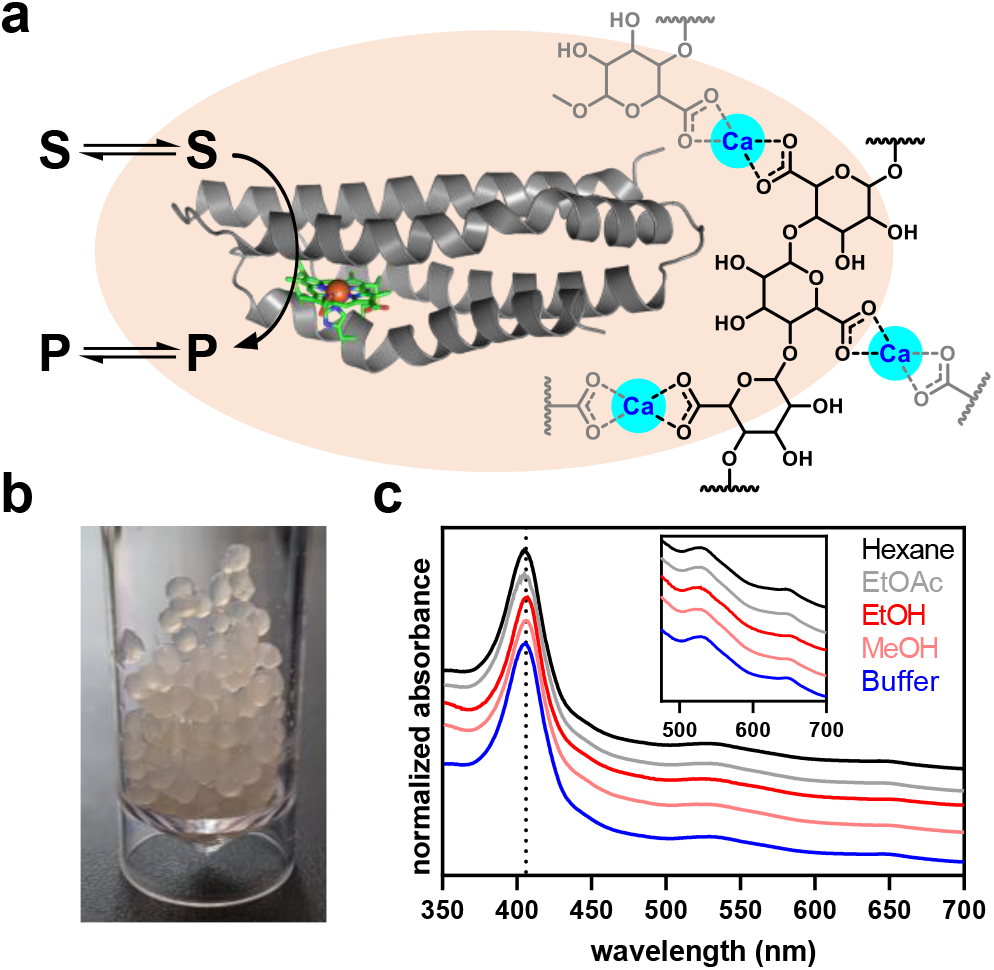
Enzyme encapsulation in calcium-alginate beads. **(a+b)** C45 is encapsulated in alginate by dropping an enzyme-alginate solution into aqueous calcium chloride. **(c)** Suspension of alginate gels in different solvents does not affect the heme absorbance spectrum.

Enzymes are readily immobilized in calcium-alginate gels by dropwise addition of a protein-alginate solution into aqueous calcium chloride (Fig. 1a).^[12]^ Instantaneous gelation occurs by virtue of ionic interactions of calcium with the alginate carboxylate groups.^[13]^ These cross-linking interactions enable enzyme immobilization but also reduce the free volume available inside the hydrogel.^[14]^ To form our C45:alginate beads, we initially determined the optimal Ca^2+^/alginate ratio for heme protein (cytochrome *c* and horseradish peroxidase) retention in aqueous solutions. As expected, leakage decreased with increasing calcium chloride concentration (Tab. S1).^[14–15]^ All further experiments were performed with beads from 300 mM Ca^2+^ and 3% alginate. At these concentrations, the density of the formed beads matched that of the aqueous phase, which provided ideal buoyancy for bead maturation. The resulting beads had an average volume of 14 μl and retained up to 33% of the tested heme-proteins, as measured by the absorbance of the heme proteins released into solution. We assessed the effects of alginate-encapsulation on C45 by creating thin C45:alginate sheets that are suitable for UV/visible spectroscopy. The characteristic Soret peak (406 nm) and Q band (530 nm and 650 nm) absorbances of the C45 heme provide insights into its structural integrity, as heme protein spectra are sensitive to changes in the local environment and the folded state of the protein. The spectra of alginate-encapsulated C45 (AE-C45) suspended in various organic solvents are virtually identical to the spectra of encapsulated or free C45 in buffer (Fig. 1), indicating that the protein structure is unaffected by the Ca^2+^:alginate matrix and effectively protected from the denaturing conditions of the neat solvents. We made similar observations with the natural cytochrome *c*, which retains its characteristic heme spectrum when encapsulated and immersed in neat organics (Fig. S1). Alginate encapsulation is thus an effective, general, and mild method for protecting the encapsulated proteins from solvent denaturation.

In *de novo* and engineered heme-containing proteins, carbene transfer reactions proceed *via* an iron(II) carbene intermediate formed by mixing ethyl diazoacetate (EDA) with ferrous heme protein.^[11]^ We therefore explored the ability of encapsulated C45 to form the ferrous and metallocarbenoid species as a precursor to probing its carbene transferase activity. On addition of the reductant sodium dithionite to C45:alginate sheets under aerobic conditions, we observed a redshift of the Soret peak to 421 nm and Q-bands at 520 nm and 550 nm, consistent with reduction to the ferrous C45 species. Notably, heme reduction inside the hydrogel was significantly slower than the near instantaneous reduction of free C45 with dithionite in solution (Fig. 2a). This indicates that diffusion of dithionite into the hydrogel is limiting the heme reduction rate, highlighting a kinetic barrier separating the alginate hydrogel and bulk solvent. After reduction to the ferrous species, we added EDA, leading to the clear formation of the metallocarbenoid intermediate, detected by a further shift of the Soret peak to 426 nm and Q-bands to 542 nm and 584 nm (Fig. 2b). The reaction of EDA with free and encapsulated C45 resulted in identical spectra (Fig. 2c), confirming the formation of the same carbene-intermediate that we have previously reported. We also note that carbene formation in the free C45 is significantly slower than heme reduction, and proceeds with a maximum rate of 0.5 s^−1^ as previously determined.^[11a]^ Thus, for slow processes such as carbene formation and transfer, equilibrium partition between solvent and bead potentially outweighs diffusion in determining the overall reaction kinetics and yield.

**Figure 2:**
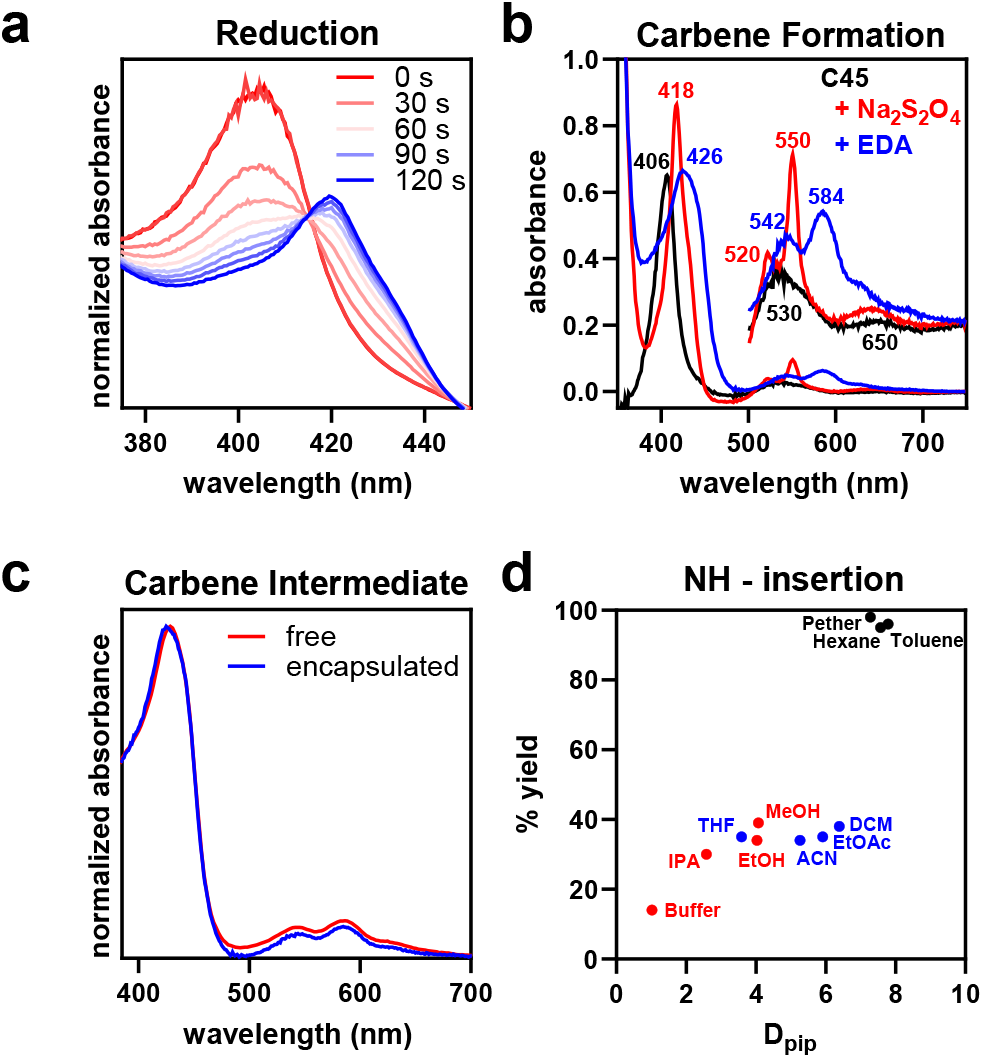
Carbene transfer activity of encapsulated C45 in organic solvent. **(a)** Slow reduction of C45 by dithionite indicates diffusion barrier across the bead surface. **(b)** Reduction and carbene formation of encapsulated C45 indicated by characteristic spectroscopic shifts. **(c)** Carbene-formation results in spectroscopically identical intermediates for free and encapsulated C45. **(d)** Product yields for N-H insertion into piperidine correlate with the substrate diffusion coefficients, indicating that yields are improved by increasing the effective substrate concentration inside the beads.

Having demonstrated that alginate-encapsulated C45 forms a metallocarbenoid intermediate, we next explored its carbene transferase activity into amine N-H bonds (Scheme 1). Using C45:alginate beads under aqueous conditions, we observed N-H insertion into piperidine when mixed with EDA and dithionite. The yield of ethyl-1-piperidine acetate (14%), as determined by HPLC (Fig. 2d, Tab. S2), was significantly lower than that observed for free C45 under the same conditions; this is consistent with previous observations of alginate-encapsulated enzymes, which often show decreased yields compared to free enzyme. However, yields could be dramatically increased by changing the solvent in which the beads were suspended, the largest gains in yield (96-98%) arising from immersion in apolar solvents such as hexane, toluene and petroleum ether. Under these conditions the performance of encapsulated C45 far exceeds that of free enzyme, which requires a 20-fold higher catalyst loading to achieve a 79% yield.^[11a]^

To determine the origin of these impressive activity gains, we examined the activity-solvent dependencies, observing an inverse relationship between reaction yield and solvent polarity (Fig S2). To further dissect this trend, we measured partition and diffusion constants of the piperidine substrate into the beads (κ_pip_ and D_pip_). We achieved this by stirring empty beads in the selected solvents containing piperidine, and subsequently measuring the decrease in piperidine concentration in the liquid phase to determine κ_pip_ and D_pip_. While both parameters exhibit some correlation to the overall N-H insertion yield, there is a near linear relationship between the D_pip_ values and yield (Fig. 2d), indicating substrate diffusion into the beads is likely rate limiting. While this analysis suggests a kinetic origin for these observed gains, we cannot exclude that equilibrium partitioning will also contribute and perhaps determine the overall reaction yield. Nevertheless, both a kinetic and equilibrium interpretation imply that a change in effective substrate concentration determines the observed yield. It should therefore be possible to exploit these trends for a desired transformation by screening and selecting the solvent system to boost product yield.

**Scheme 1.**
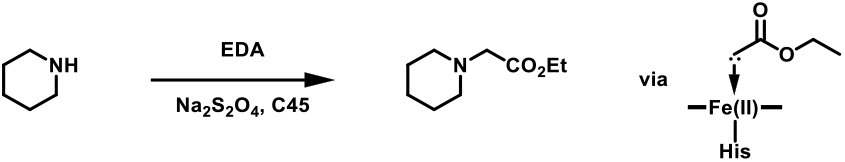
C45 mediated NH-insertion via carbene-transfer

We further probed the effect of partition on the peroxidase activity of C45. Based on our previous observations, it was expected that immersing C45:alginate beads in organic solvents would increase the apparent peroxide concentration in the bead. Given that high concentrations of peroxide often have deleterious effects on enzyme activity, encapsulation should therefore result in significant activity losses. We analyzed the peroxidase activity of C45 encapsulated in alginate beads by monitoring the oxidation of guaiacol by H_2_O_2_ (Fig. 3 and Scheme 2). C45:alginate beads in buffer retained peroxidase activity with a *k*_cat_ of 170 s^−1^ under limiting guaiacol concentration (1 mM), which is only 11-fold lower than observed for free C45. The apparent *K*_m_ decreased 4.5-fold upon encapsulation, signaling an increased apparent peroxide concentration inside the beads, which likely also causes the apparent drop in *k*_cat_ through partial enzyme inactivation. We observed analogous effects when using ABTS as the electron donor in the peroxidase reaction, with a diminished *k*_cat_ and elevated *K*_M_ relative to those of free C45 under the same reaction conditions. Suspending the beads in organic solvents drastically decreased C45’s peroxidase activity, which likely signals an even higher effective peroxide concentration inside the beads due to partitioning between the organic and aqueous phases. Substituting C45 for horseradish peroxidase yielded similar results, with HRP:alginate beads in buffer retaining some activity for ABTS oxidation after encapsulation (Fig. S3). These effects on peroxidase activity support our model, signaling an increased apparent substrate concentration by partitioning. This may explain the activity losses observed for other systems,^[9d, 9e]^ and suggest opportunities to exploit substrate partitioning to boost enzyme activity in specific cases.

**Figure 3:**
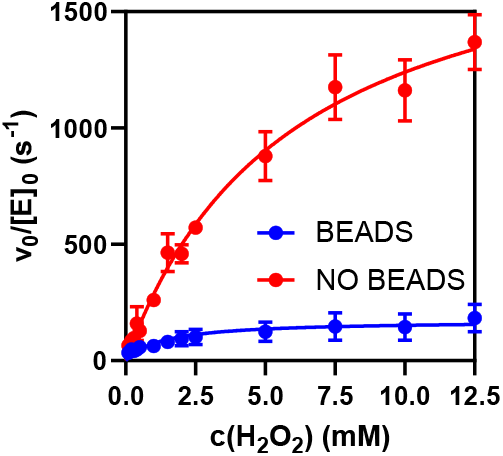
Peroxidase activity. C45 encapsulated in calcium-alginate beads retained activity for guiacol oxidation. Activity is decreased, likely due to non-beneficial partitioning of peroxide into the beads.

**Scheme 2.**
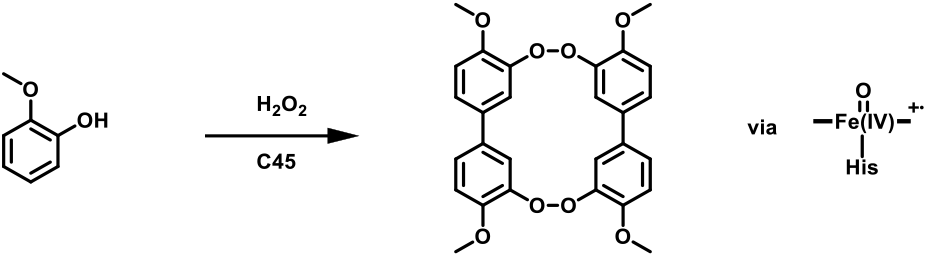
C45 mediated oxidation of guaiacol *via* Compound I

Enzyme recyclability is highly desirable from an industrial perspective, and alginate-beads can be readily recovered and reused. To test the reusability of our encapsulated C45 in buffer, acetonitrile and tetrahydrofuran, beads were first used to catalyze the cyclopropanation of styrene, (Fig. 4, Tab. S3 and Scheme 3). After removal of the reaction mixture, the same beads were subsequently used to catalyze carbene N-H insertion into piperidine. The cyclopropanation proceeded with moderately high yields (62-79%) and comparable enantioselectivity to free C45 (86-99%). This stereoselectivity further indicates that C45 remains active and folded after encapsulation. After isolating and washing the beads, we further tested their N-H insertion activity against piperidine, observing between 30-65% residual activity depending on the solvent used for the reaction. We did not detect any cyclopropanation product in the HPLC chromatograms of the N-H insertion reactions. The absence of cross contamination with residual reagents as well as formation of different product during the second reaction clearly confirms that the encapsulated C45 can be recovered and remains active towards carbene transferase chemistry in successive reactions.

**Figure 4:**
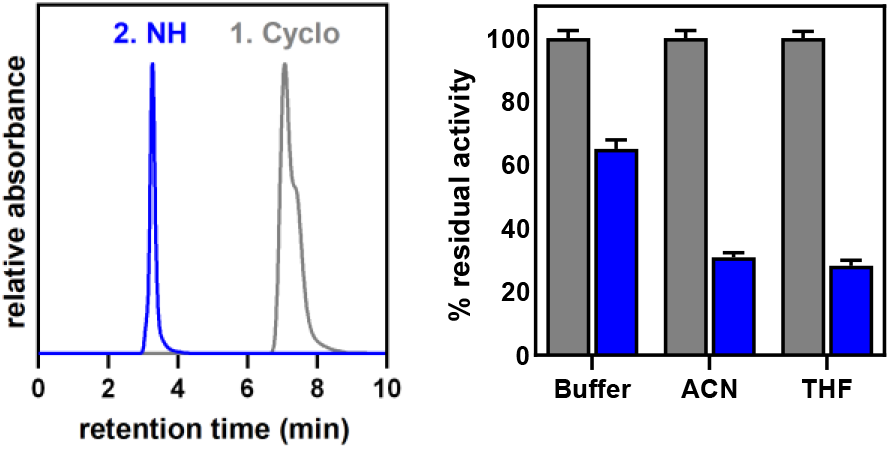
Encapsulated enzymes can be readily recycled. Encapsulated C45 has been employed to sequentially promote cyclopropanation (1. Cyclo, black) followed by NH-insertion reaction (2. NH, blue) with some losses in activity.

**Scheme 3.**
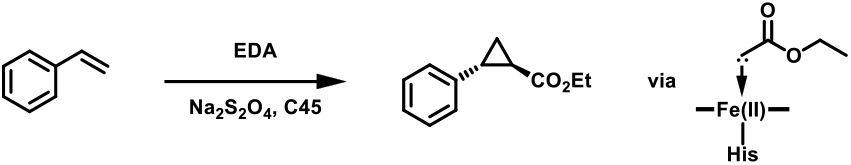
C45 mediated cyclopropanation via carbene-transfer

We have demonstrated here that enzyme encapsulation in alginate beads enables non-aqueous biocatalysis under solvent conditions that are detrimental to the free enzyme. The beads introduce a diffusion barrier, that is rate limiting for peroxidase and carbene transferase activities. The effective substrate concentration inside the bead is key to controlling activity and can be manipulated by screening and selecting the optimal solvent system for the desired reaction. Recoverability and recyclability are added application benefits of such alginate-encapsulation. The immobilization of *de novo*, engineered, and natural enzymes in alginate beads is therefore a versatile method and this work provides a rational basis to tune solvent systems to maximize product yields.

## ASSOCIATED CONTENT

### Supporting Information

Supporting Information contain complete experimental procedures and additional kinetic data, including Table S1-S3 and Figures S1-S3.

## AUTHOR INFORMATION

### Author Contributions

R.S. and H.A.B. contributed equally. R.S., H.A.B., and J.L.R.A. designed the experiments. R.S. and H.A.B. performed the experiments. R.S., H.A.B., and J.L.R.A. wrote the manuscript with input from A.J.M.

### Notes

The authors declare no conflict of interest.

## ACKNOWLEDGMENTS

This work was supported by the Biological and Biotechnological Sciences Research Council (BBI014063/1, BB/R016445/1, and BB/M025624/1), the Engineering and Physical Sciences Research Council (EP/R026939/1, EP/R027129/1 and EP/M022609/1) and the Bristol Centre for Functional Nanomaterials (Engineering and Physical Sciences Research Council Doctoral Training Centre Grant EP/G036780/1) through a studentship for RS. HAB thanks the Swiss National Science Foundation for a Postdoc.Mobility fellowship.

