## Supporting Information for "Beneficial substrate partitioning boosts non-aqueous catalysis in de novo enzyme-alginate beads"

### 1. Supporting Tables

**Table S1: Alginate bead volumes and enzyme retention.**

| 3% (w/v) sodium alginate | $CaCl_2$ | | | | |
|---|---|---|---|---|---|
| | **0.1 M** | **0.2 M** | **0.3 M** | **0.4 M** | **0.5 M** |
| **Particle volume – hydrated** | 14.1 µl | 14.1 µl | 14.1 µl | 14.1 µl | 14.1 µl |
| **Particle volume – dehydrated** | 3.0 µl | 4.2 µl | 4.9 µl | 7.2 µl | 11.5 µl |
| **Water volume** | 11.1 µl | 9.9 µl | 9.2 µl | 6.9 µl | 2.6 µl |
| **Water content** | 78% | 70% | 66% | 49% | 19% |
| **HRP retention** | 11% | 15% | 24% | 28% | 34% |
| **Cytochrome *c* retention** | 19% | 24% | 33% | 40% | 46% |

**Table S2: N-H insertion reaction yields in different solvents.**

| Solvent | % Yield | polarity index | $K_{piperidine}$ | $D_{piperidine}$ |
|---|---|---|---|---|
| **Buffer** | 14 ± 2 | 10.2 | 0.08 | 1.02 |
| **MeOH** | 39 ± 2 | 5.1 | 0.50 | 4.07 |
| **EtOH** | 34 ± 1 | 4.3 | 0.48 | 4.03 |
| **IPA** | 30 ± 3 | 3.9 | 0.20 | 2.58 |
| **ACN** | 34 ± 1 | 5.8 | 1.10 | 5.26 |
| **EtOAc** | 35 ± 1 | 4.4 | 1.78 | 5.91 |
| **THF** | 35 ± 1 | 4 | 0.37 | 3.59 |
| **DCM** | 38 ± 2 | 3.1 | 2.59 | 6.39 |
| **Hexane** | 95 ± 3 | 0.1 | 7.21 | 7.57 |
| **Pether** | 98 ± 2 | 0.1 | 5.57 | 7.28 |
| **Toluene** | 96 ± 2 | 2.4 | 8.85 | 7.79 |
| **free enzyme [a]** | 79 ± 12 a | n. A. | n. A. | n. A. |

[a] Reaction of the free enzyme in CHES buffer, 20-fold higher catalyst loading compared to the reactions inside the beads. Value from Stenner et al., PNAS 2020.[1]

**Table S3: Sequential cyclopropanation and N-H insertion with recovered beads [a]**

| Solvent | Cyclopropanation | | | N-H insertion |
|---|---|---|---|---|
| | **% Yield(R,R)** | **% Yield(S,S)** | **ee(R,R)** | **% Yield** |
| **Buffer** | 63.5 ± 1.5 | 4.73 ± 0.78 | 86% | 9.0 ± 0.4 |
| **ACN** | 77.6 ± 1.8 | 0.12 ± 0.05 | ≥99% | 10.4 ± 0.5 |
| **THF** | 68.0 ± 1.4 | 0.14 ± 0.01 | ≥99% | 9.9 ± 0.6 |

[a] Alginate encapsulated C45 catalyzed cyclopropanation reaction between EDA and styrene followed by the recovery and reuse of the same beads in the N-H insertion reaction between EDA and piperidine.

#### 2. Supporting Figures

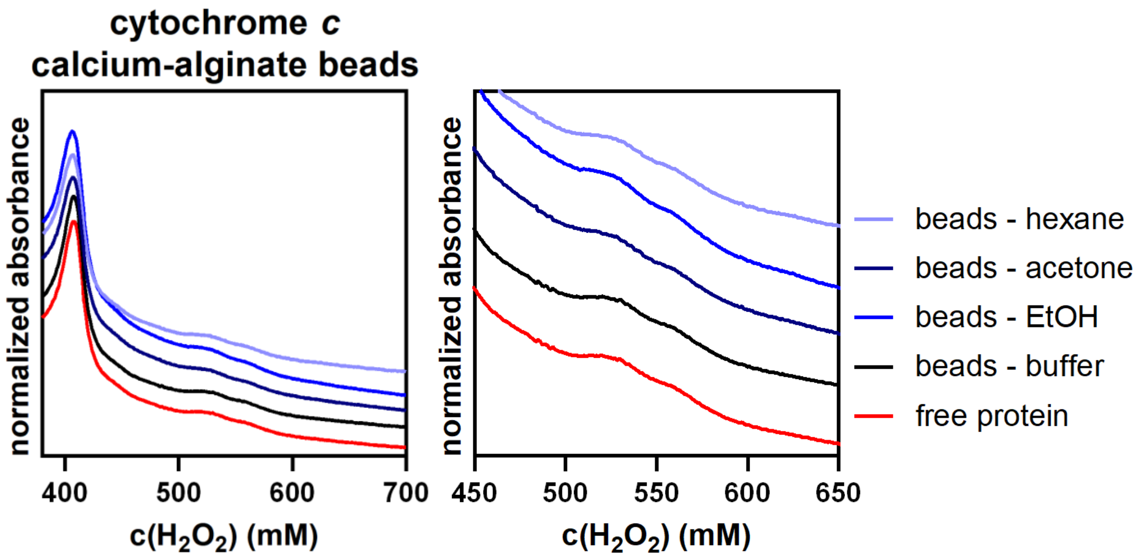

**Figure S1: UV-VIS spectra of cytochrome c in beads.** Spectra of encapsulated cytochrome *c* in various solvents are almost identical to the spectrum of free protein in buffer.

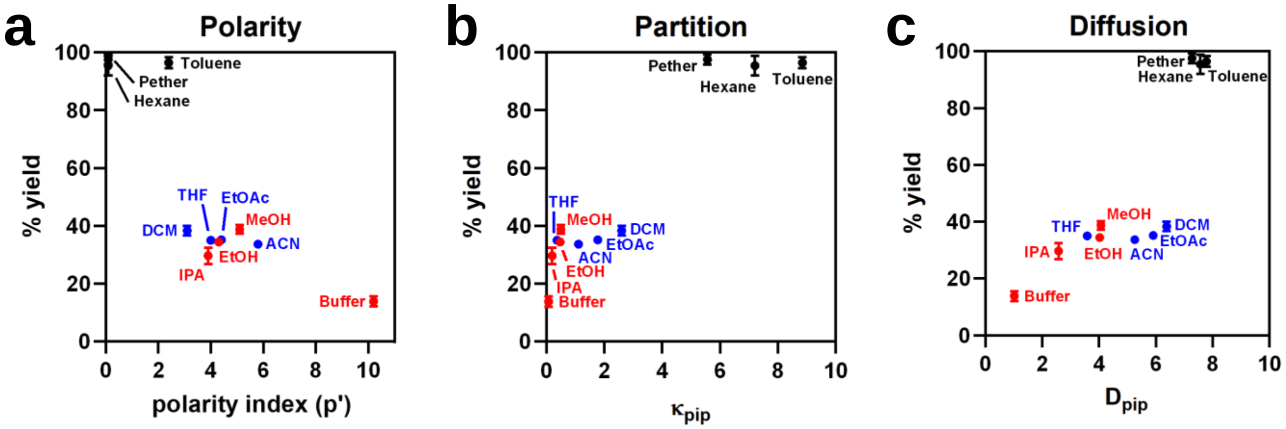

**Figure S2: Partition versus diffusion. (a)** Polarity index (p'),[2] **(b)** diffusion ($D_{pip}$) and **(c)** partition ($\kappa_{pip}$) coefficients of the piperidine substrate correlate well with the N-H insertion yields of alginate-encapsulated C45. The slow reduction of heme inside the beads suggests that both the ensuing carbene formation and N-H insertion will be limited by substrate diffusion into the beads. Our data indicate a kinetic effect, but we cannot exclude that equilibrium partition will be determining the overall yield. Nonetheless, both a kinetic and equilibrium interpretation imply that increased apparent substrate concentration results in the observed superior yields.

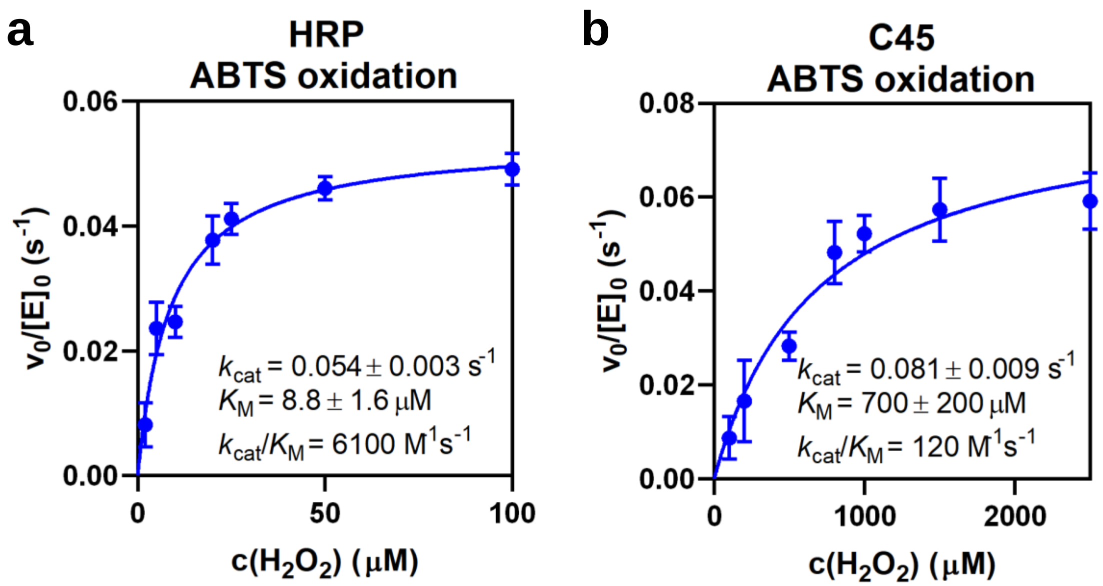

**Figure S3: Peroxidase activity.** Alginate encapsulation is mild and general method for enzyme immobilization. Both C45 **(a)** and HRP **(b)** remain active inside the beads when activity is measured ABTS (1 mM). As expected, peroxidase activities are decreased compared to the free enzyme,[3] which is likely due to an increase in apparent peroxidase concentration inside the bead that leads to enzymes degradation.

### 3. Materials and Methods

Chemicals were purchased from either Sigma or Fisher Scientific. All anaerobic assays were performed in a glove box under $N_2$ ([$O_2$] <5 p.p.m.; *Belle Technology*). LC-MS mass spectra were acquired using positive electron-spray-ionization (ESI) mass spectrometry (*Waters* Xevo G2-XS QTof) attached to a C8 reverse column (*Grace Vydac*, 100 x 2.1 mm, 5 µm). UV-Vis spectroscopy was performed with an Agilent Cary-60 UV-visible spectrophotometer using a 1 cm pathlength quartz cuvette. HPLC chromatograms were acquired using an Agilent waters HPLC and three separate columns were employed for quantification: a C18 HPLC reverse phase column (*Phenomenex*, 150 x 15 mm, 5 µm) and a CHIROBIOTIC® V chiral-HPLC column (*Astec*, 25 x 2.1 mm, 5 µm).

#### 3.1 Protein production and purification

C45 was expressed and purified as described previously.[3a] Briefly, *E. coli* T7 Express competent cells (*NEB*) were transformed with plasmids harboring C45 and pEC86+ to assist *c*-type heme maturation. Cells were grown overnight at 37 °C and 180 rpm. 1 L cultures were inoculated with this overnight culture and grow to an $OD_{600}$ of 0.6-0.8 at 37 °C and 180 rpm. Protein production was induced with 1 mM IPTG. After four hours at 37 °C and 180 rpm, cultures were harvested by centrifugation (4000 rpm, 4 °C, 20 mins) and the resultant cell pellets were isolated from the supernatant, resuspended in 10 ml lysis buffer (300 mM NaCl, 50 mM sodium phosphate, 20 mM imidazole, pH 8) and stored at -20 °C until purification. The lysis suspensions were thawn, sonicated and the lysate cleared by centrifugation (16000 rpm, 4 °C, 30 mins) prior to affinity purification on a Ni-NTA column. The His-tag was cleaved by TEV protease and uncleaved protein was removed by running the protein through a Ni-NTA column. The C45 concentration was determined using the extinction coefficients of the ferric Soret-peak ($\varepsilon_{406nm}$ = 147,000 $M^{-1}cm^{-1}$).[3a] Pure protein was obtained after subsequent size-exclusion purification over a S75 gel filtration column in CHES buffer (20 mM CHES, 100 mM KCl, pH 8.6) and stored at -80 °C.

#### 3.2 Alginate encapsulation

A solution of 3% (w/v) sodium alginate was prepared by mixing 3 g of sodium alginate with 100 ml of CHES buffer (20 mM CHES, 100 mM KCl, pH 8.6). The mixture was left stirring for 1 h to ensure complete dissolution and homogeneity. 1 ml of 100 µM C45 final concentration 10 µM) were added to 9 ml of 3% (w/v) sodium alginate and adequately mixed to ensure homogeneity. To prepare thin alginate-sheets for UV-Vis spectroscopy, a thin film of 0.2 M $CaCl_2$ was prepared in a petri dish and the alginate-enzyme mixture was slowly pipetted over the surface. Sheets were cured for 30 minutes before the residual $CaCl_2$ was discarded and the sheets were air-dried for 30 min and transferred into a cuvette for UV-VIS analysis. To prepare alginate-beads for biocatalytic assays, the alginate-enzyme mixture was added dropwise from an elevation of approximately 30 cm into a conical flask equipped with a stirrer bar and containing an excess of 0.2 M $CaCl_2$. The beads formed spontaneously and were cured for 30 min under stirring before being collected by vacuum filtration. The beads were air-dried for 30 minutes before being stored at 4 °C.

#### 3.3 Chemophysical characterization of alginate beads

##### 3.3.1 Volumetric analysis

For the volumetric analysis, alginate beads without protein were prepared as described above, by dropping a solution of sodium alginate into $CaCl_2$ solutions of varying concentrations (0.1 – 0.5M). The beads were matured for 1 h in the respective $CaCl_2$ solution. Beads were collected by vacuum filtration and subsequently weighted and measured. Beads were incubated at 37 °C for 3 h. After 3 hours, the beads were weighted and measured again to determine the water content of each bead.

##### 3.3.2 Retention analysis

10 mg of horseradish peroxidase or cytochrome c were dissolved in 10 ml of a sodium alginate solution (3% (w/v) sodium alginate, 20 mM CHES, 100 mM KCl, pH 8.6) and mixed on a roller to achieve homogeneity. Alginate beads were prepared as described above, by dropping a solution of sodium alginate into $CaCl_2$ solutions of varying concentrations (0.1 – 0.5 M). The beads were matured for 1 h in the respective $CaCl_2$ solution before being collected via vacuum filtration and resuspended in 50 ml of CHES buffer (20 mM CHES, 100 KCl, pH 8.6). The samples were mixed on a roller overnight. The concentration of each protein leaked into the aqueous phase was determined from the absorbance of the Soret peak (HRP, $\varepsilon_{403nm}$ = 103,840 $M^{-1}cm^{-1}$; cytochrome c, $\varepsilon_{409nm}$ = 98,160 $M^{-1}cm^{-1}$).

##### 3.3.3 Bead-solvent partition and diffusion coefficients

Partition coefficient, alginate bead porosity and diffusion coefficients were determined following published procedures.[4]

**Partition coefficient.** The partition coefficient $\kappa_{pip}$, is defined by Equation 1, where $[pip]_{bead}$ and $[pip]_{bulk}$ are the concentrations of a piperidine in the alginate-bead and in the bulk solution, respectively. C18-HPLC was used to determine the $\kappa_{pip}$ in various solvents using alginate beads without enzyme. An external calibration for piperidine was performed by injecting various concentrations of piperidine and recording the peak height in the chromatogram at 254 nm (piperidine, concentration range 500 µM – 10 mM). To a 2 ml screw-top vial was added 990 µl of the respective solvent, 10 µl of a 3 M stock of piperidine in EtOH and three alginate beads. The final concentration of piperidine was 10 mM. The vials were mixed on a roller for 2 hours before the beads were removed from the vials prior to loading the solvents onto the HPLC. The concentration of the substrate remaining in the bulk solution was calculated from the peak height. The partition coefficient was calculated according to Equation 1, after determining the $[pip]_{bead}$ from the loss of piperidine in the bulk solvent.

$$\kappa_{\mathrm{pip}} = \frac{[\mathrm{pip}]_{\mathrm{bead}}}{[\mathrm{pip}]_{\mathrm{bulk}}} \qquad \text{Equation 1}$$

**Alginate porosity.** The porosity (ε) of the alginate beads was determined according to Equation 2. Alginate beads without enzyme were suspended in 50 ml of 100 mM indigo carmine and allowed to stir for 20 minutes. After two hours the beads were collected via vacuum filtration and the concentration of indigo carmine remaining in the 50 ml solution ($C_1$) was determined via UV-VIS spectroscopy ($\varepsilon_{667nm}$ = 34,145 $M^{-1}cm^{-1}$). The collected beads were then resuspended in 50 ml ($V_b$) of 17 mM indigo carmine ($C_2$) and left mixing for 20 min, before the beads were collected via vacuum filtration and the concentration of indigo carmine in the second solution was determined using UV-VIS spectroscopy (C). The porosity of the alginate beads was calculated using Equation 2:

$$\varepsilon = \frac{3V_b(C-C_2)}{4\pi r^3 N(C_1-C)} = \frac{V_b(C-C_2)}{V_p(C_1-C)} \qquad \text{Equation 2}$$

where N is the number of beads in the solution, $V_p$ is total bead volume and r the radius of a single alginate bead. For alginate beads prepared from 0.3M $CaCl_2$ and 3% (w/v) sodium alginate the porosity was 0.89.

**Diffusion coefficient**. The diffusion coefficients of piperidine into the beads in various solvents was determined according to the non-steady state Equation 3 that assumes that all beads are perfectly spherical and Fickian diffusion is the only mechanism of substrate transport in the radial direction.

$$S_b(t) = \frac{\alpha S_b(0)}{1+\alpha}\left[1 + 6(1+\alpha)\sum_{n=1}^{\infty}\frac{e^{\left(\frac{-D_{pip}q_n^2 t}{r^2}\right)}}{9+9\alpha+q_n^2\alpha^2}\right] \quad \text{Equation 3}$$

Where $S_b(t)$ is the concentration of substrate in the bulk at time t, $S_b(0)$ the concentration of substrate in the bulk at time 0, r the radius of the alginate beads, and $D_{pip}$ the diffusion coefficient. To solve Equation 3, the values of α and $q_n$ were determined using Equation 4 and Equation 5. α, and thus the diffusion coefficient, was calculated based on the partition coefficient $\kappa_{pip}$ determined above, with $V_b$ being the volume of the bulk solution and $V_p$ the total volume of the beads. The non-positive roots for Equation 5, and the value of $q_n$ was solved using the Secant method.

$$\alpha = \frac{V_b}{V_p \kappa_{pip}} \quad \text{Equation 4}$$

$$\tan(q_n) = \frac{3q_n}{3+\alpha q_n} \quad \text{Equation 5}$$

Since the effective diffusion ($D_{pip,eff}$) and $D_{pip}$ are related through porosity (ε), $D_{eff}$ can be calculated from Equation 6:

$$D_{pip,eff} = \varepsilon D_{pip} \quad \text{Equation 6}$$

#### 3.4 Kinetic assays

##### 3.4.1 Guaiacol oxidation

Oxidation of guaiacol was assayed at a final enzyme concentration of 28 nM in a total volume of 1 ml. To that end, one alginate bead (14 µl, c(C45) = 2 µM) was added to solutions containing $H_2O_2$ and the substrate and the reaction proceeded under constant stirring. Kinetic assays were performed at a concentration of 1 mM of guaiacol, while the concentration of peroxide varied between 250 and 2500 µM. Product formation was monitored by absorbance using a $\Delta\varepsilon_{470nm}$ = 26,600 $M^{-1}cm^{-1}$. All reactions were repeated in triplicates. Initial rates were calculated from the slope fitted to the Michaelis-Menten equation.

##### 3.4.2 UV-vis spectra of the metallocarbenoid intermediate

To a 1 cm pathlength UV-VIS quartz cuvette was added a thin-sheet of alginate-encapsulated C45. 1 ml CHES buffer (20 mM CHES, 100 mM KCl, pH 8.6) was added to the cuvette and a UV-VIS spectrum was recorded. An initial baseline was acquired using an alginate sheet without any enzyme encapsulated within it. The cuvette was subsequently sealed with a silicon septum and flushed with $N_2$ from a nitrogen cylinder. The reduced spectrum was acquired by adding 10 µl of a 100 mM sodium alginate solution (in de-ionized water) into the cuvette containing the solvent/alginate-encapsulated C45 sheet via an airtight syringe. A UV-VIS spectrum was then recorded every 30 s. After sufficient reduction could be detected, 10 µl of a stock solution of ethyl diazoacetate (EDA, 400 mM in EtOH) was added into the sealed cuvette via an airtight syringe and UV-VIS spectra were every 30 s for 10 min.

##### 3.4.3 Piperidine N-H insertion.

All reactions were prepared in an anaerobic glove box. To 970 µl of solvent (CHES buffer, MeOH, EtOH, isopropanol, acetone, acetonitrile, ethyl acetate, dichloromethane, THF, n-hexane, petrol ether, toluene) in a 1.5 ml glass vial was added 3 alginate-enzyme (10 µM, 14 µl) beads and 10 µl of $Na_2S_2O_4$ (1M stock; deionized water) and the mixture was left to stir for 1 minute. 10 µl of a piperidine (3M stock; EtOH) was added and the reaction left to mix for 30 seconds. After 30 seconds 10 µl of EDA (1M stock; EtOH) was added into the reaction vials to initiate the reaction. Final concentrations were 30 μM enzyme, 10 mM sodium dithionite, 10 mM diazo compound, and 30 mM piperidine. Once mixed, the reactions were mixed on a roller at room temperature. After 2 hours, the reaction vials were unscrewed, the beads were removed, and the mixture was transferred to a 15 ml falcon tube containing 2 ml of a 1:1 water:ethyl acetate mixture, vortexed and then centrifuged (14,500 rpm; 2 minutes). The organic layer was isolated and subsequently analyzed by chiral-HPLC and LC-MS. The product yields and total turnover numbers (TTN; concentration of product formed/concentration of enzyme) where calculated via an external calibration with commercial ethyl-1-piperidineacetate.

##### 3.4.4 Recycling of beads used for cyclopropanation in N-H insertion assay

Recoverability assays were conducted inside 1.5 mL screw top vials sealed with a silicone-septum containing cap. Assays were prepared in an anaerobic glove box in either CHES buffer (100 mM KCl, 20 mM CHES, pH 8.6) or the specified organic solvent (acetonitrile or THF). The final reaction volumes for all assays were 1 ml. To 970 μl of solvent in a 1.5 ml glass vial was added 3 alginate-enzyme (10 μM, 14 μl) beads and 10 μl of $Na_2S_2O_4$ (1M stock; deionized water) and the mixture was left to stir for 1 minute. 10 μl styrene (3M stock; EtOH) was added and the reaction left to mix for 30 seconds. After 30 seconds 10 μl of EDA (1M stock; EtOH) was added into the reaction vials to initiate the reaction. Final concentrations were 30 μM enzyme, 10 mM sodium dithionite, 10 mM diazo compound, and 30 mM styrene. Once mixed, the reactions were stirred on a roller at room temperature. After 2 hours, the reaction was quenched by the addition of 20 μl of 3M HCl. The vials were unscrewed, the beads were removed, and the resultant mixture was transferred to a 15 ml falcon tube containing 1 ml of ethyl acetate, vortexed and then centrifuged (14,500 rpm; 2 minutes). The organic layer was then either i) analyzed via chiral-HPLC, or ii) collected and then slowly added to a 15 ml falcon tube containing 1 ml of an aqueous 3M NaOH solution. The resulting biphasic mixture was left to stir, at room temperature, for 2 hours; the progress of the hydrolysis reaction was monitored by TLC (7:3 ethyl acetate:hexane, 254 nm). After 2 hours, the aqueous layer was isolated analyzed by chiral-HPLC and LC-MS. The product yields and enantiomeric excesses where calculated via an external calibration with commercial ethyl 2-phenylcyclopropane-1-carboxylate and 2-phenylcyclopropane-1-carboxylic acid. After each reaction, the sealed vials were transported into an anaerobic glove box the lids were removed and the product mixture extracted was decanted into an Eppendorf for analysis. The beads were washed twice with the respective solvent before assaying N-H insertion activity as described above.

#### 3.5 Product characterization

##### 3.5.1 Product characterization by reverse phase and chiral HPLC

N-H insertion and cyclopropanation reactions were analyzed by high performance liquid chromatography. Three separate columns were employed for quantification: two chiral columns and a reverse phase C18 column. An Astec CHIROBIOTIC® V (25.0 mm x 2.1 mm, 5 μm) chiral-HPLC column and a CYCLOBOND® I (21.0 mm x 2.1 mm, 5 μm) chiral-HPLC column was used to analytically quantify the cyclopropanation and N-H insertion assays, employing an isocratic mobile phase (100% CH3CN: 0.1% v/v TFA: 0.1% v/v: Et3N; 0.1 ml $min^{-1}$ flow rate and 2 μl injection volume for the cyclopropanation assays; 0.2 ml $min^{-1}$ flow rate and 5 μl injection volume for the N-H insertion assays) for the CHIROBIOTIC® V column and an isocratic mobile phase (95:5 v/v n-hexane:isopropanol; 0.4 ml $min^{-1}$ flow rate and 10 μl injection volume for the cyclopropanation and N-H insertion assays) for the CYCLOBOND® I column. All elution traces were monitored spectroscopically at 245, 254 and 280 nm. The chiral-HPLC column allowed the retention times of the starting materials and the reaction products to be determined, and to quantify, where necessary, the enantioselectivity of a given reaction. A C18 HPLC reverse phase column (Phenomenex, 150 mm x 15 mm, 5 μm) was also employing for quantitative analysis of the chemophysical properties of the alginate beads; either a gradient mobile phase (70:30% H2O:$CH_3CN$ to 10:90% H2O:$CH_3CN$; 2 ml $min^{-1}$ flow rate and 20 μl injection volume) or an isocratic mobile phase (100% CH3CN: 0.1% v/v TFA: 0.1% v/v: Et3N;

2 mL min-1 flow rate and 20 µl injection volume) was employed. All elution traces were monitored spectroscopically at 245, 254, 265 and 280 nm.

##### 3.5.2 Calibrations curves

Calibrations curves of the reaction products were obtained as described previously.[1] Briefly, the conditions and mobile phases employed in the activity analysis were employed to prepare each substrate at multiple concentrations and the peak height in the chromatogram was recorded as a function of concentration.

##### 3.5.3 Liquid chromatography-Mass spectrometry

Liquid chromatography-Mass spectrometry was employed to identify the products formed in each assay. A C8 reverse column (Grace Vydac, 100 mm x 21 mm, 5 µm) was used with a 10-minute gradient mobile phase (95:5% H2O:CH3CN to 10:90% H2O:CH3CN; 0.1% v/v formic acid, 0.25 ml min$^{-1}$) for sample analysis. 20 µl sample were injected onto the column. After eluting from the column, the mobile phase entered an isocratic solvent chamber where a 1:100 dilution was performed prior to injecting the sample into a positive electron-spray-ionization mass spectrometer (Waters Xevo G2-XS QTof). Masses were screened across a m/z range of 70-350. Retention times and fragmentation patterns were determined for commercially available standards.
